# APOBEC3C tandem domain proteins create super restriction factors against HIV-1

**DOI:** 10.1101/2020.03.27.012963

**Authors:** Mollie M. McDonnell, Kate H.D. Crawford, Adam S. Dingens, Jesse D. Bloom, Michael Emerman

**Author notes:** Corresponding author. Michael Emerman, 1100 Fairview Avenue N., C2-023, Seattle WA, USA 98109.

## Abstract

Humans encode proteins, called restriction factors, that inhibit replication of viruses like HIV-1. One family of antiviral proteins, *apolipoprotein B mRNA-editing enzyme catalytic polypeptide-like 3 (*APOBEC3, shortened to A3) acts by deaminating cytidines to uridines during the reverse transcription reaction of HIV-1. The *A3* locus encodes seven genes, named *A3A*-*A3H*. These genes either have one or two cytidine deaminase domains and several of these A3s potently restrict HIV-1. A3C, which has only a single cytidine deaminase domain, however, inhibits HIV-1 only very weakly. We tested novel double domain protein combinations by genetically linking two *A3C* genes to make a synthetic tandem domain protein. This protein created a “super restriction factor” that had more potent antiviral activity than the native A3C protein, which correlated with increased packaging into virions. Furthermore, disabling one of the active sites of the synthetic tandem domain protein results in an even greater increase in the antiviral activity—recapitulating a similar evolution seen in A3F and A3G (double domain A3s that only use a single catalytically active deaminase domain). These A3C tandem domain proteins do not have an increase in mutational activity, but instead inhibit formation of reverse transcription products which correlates with their ability to form large higher order complexes in cells. Finally, the A3C-A3C super restriction factor largely escaped antagonism by the HIV-1 viral protein, Vif.

**Importance:** As a part of the innate immune system, humans encode proteins that inhibit viruses like HIV-1. These broadly acting antiviral proteins do not protect humans from viral infections because viruses encode proteins that antagonize the host antiviral proteins to evade the innate immune system. One such example of a host antiviral protein is APOBEC3C (A3C), which weakly inhibits HIV-1. Here, we show that we can improve the antiviral activity of A3C by duplicating the DNA sequence to create a synthetic tandem domain, and furthermore, are relatively resistant to the viral antagonist, Vif. Together, these data give insights about how nature has evolved a defense against viral pathogens like HIV.

## Introduction

The *APOBEC3 (A3)* gene locus in primates encodes cytidine deaminase proteins that inhibit endogenous retroelements, like LINE-1, and retroviruses, such as HIV-1, among other viruses including hepatitis B virus, human papillomavirus, and some herpes viruses (1–4). This *A3* gene locus has expanded in primates to give rise to the seven members of the *A3* family, named *A3A* to *A3H* (5). For A3 proteins to inhibit HIV-1 replication, they must be packaged into budding virions and brought to the target cell, where they extensively deaminate cytidines on the negative strand of ssDNA to uridines during reverse transcription (1, 2). The resultant G-to-A hypermutation on the positive strand renders the provirus non-functional. The most potent naturally found A3, A3G, can mutate up to 10% of the guanines on a single provirus (6). In addition to extensive hypermutation, some A3s can also inhibit reverse transcription via deamination-independent mechanisms as demonstrated by the presence of truncated cDNA products in the presence of A3s (1, 7–10). Recent studies of A3G show that this non-enzymatic mechanism functions by binding and sterically hindering the enzymatic activity of reverse transcriptase (11).

There are four human A3s that have endogenous antiviral activity against HIV-1 in T cells: A3D, A3F, A3G, and A3H with A3G being the most potent (12). In order to replicate in the presence of these A3s, HIV-1 and other lentiviruses encode a protein, Vif, that targets A3s for proteasomal degradation. Vif has evolved three separate interfaces in order to degrade A3s: one binding A3G, another one A3H, and a third able to recruit A3C/A3D/A3F (2). Moreover, Vif also binds to the host factor CBF-β to help recruit an E3 ubiquitin ligase complex composed of CUL5, ELOB, ELOC, RBX2, and ARIH2 proteins that mediate polyubiquitination and rapid degradation of the A3s through the proteasome (13, 14).

In addition to the multiple *A3* genes in humans, there is additional diversity in the *A3* locus in the form of human polymorphisms that affect the antiviral activity against HIV-1. For instance, there are at least twelve haplotypes of A3H that vary in their ability to restrict HIV-1 (15, 16). In addition, the most common form of human A3C, which encodes for a serine at position 188, has little, if any, activity against HIV-1. However, about 10% of African individuals carry a polymorphism in A3C that encodes an isoleucine at position 188. This one amino acid change from a serine to an isoleucine is correlated with increased antiviral activity against HIV-1 resulting from an increased ability of A3C to form dimers and increased cytidine deaminase activity *in vitro* (17). Furthermore, chimpanzee and gorilla A3Cs have different amino acids at position 115 than humans and introducing these amino acids into human A3C also increase dimerization and cytidine deaminase activity (18). Nonetheless, even with these mutations, A3C is less active against HIV-1 compared to many of the other A3 proteins (19).

Each *A3* gene can be classified according to the presence of a zinc-coordinating domain motif: A3Z1, A3Z2, or A3Z3 (20). The evolutionary history of the *A3* locus is characterized by duplication, recombination, and deletion events; in the last 50 million years, three duplication events have occurred in the A3Z1 and A3Z2 sub-families, but not in the A3Z3 (5). These fusion and recombination events gave rise to the seven *A3*s found in primates that include the three single deaminase domain *A3s* (*A3A, A3C*, and *A3H*) and four double deaminase domain *A3s* (*A3B, A3D, A3F*, and *A3G*).

Super restriction factors are defined as evolution-guided variants of natural restriction factors with improved properties based on previous work done on the restriction factor MXA (21). Super restriction factors provide a unique tool to study how restriction factors work and the evolutionary compromises for paths that have yet to be sampled in nature. Because the most active A3s are double domain proteins, we examined the hypothesis that super restrictor factors could be made by duplicating the poorly active single domain A3C protein into a synthetic tandem domain protein. We did this for both the common form of human A3C (A3C_S188_) and the more active variant of A3C (A3C_I188_). Remarkably, we found that all A3C tandem domain variants have greater antiviral activity relative to their single domain counterparts. We found that the tandem deaminase A3C variants are packaged into virions at higher levels than their single domain counterparts, likely explaining the majority of the increase in antiviral activity. In the natural double domain APOBEC3 proteins, only the C-terminal cytidine deaminase active sites are used for hypermutation (22, 23). Here, we show that mutation of the cytidine deaminase active site in the C-terminal domain of A3C_S188_-A3C_S188_ results in even greater antiviral activity than the same protein with two active sites. This increase in antiviral activity is correlated with elevated packaging into virions and is largely independent of increased mutational load. Instead, there were lower amounts of reverse transcriptase products in cells infected with virus produced in the presence of these A3C tandem domain proteins. This inhibition of late reverse transcription products is also correlated with the formation of large higher-order complexes in the A3C-A3C variants compared to their single domain counterparts. Finally, we show that the A3C tandem domain proteins are largely resistant to antagonism by HIV-1 Vif. These A3C-A3C super restriction factors provide a unique tool to understand evolutionary paths of antiviral genes and their antagonism by viral proteins.

## Results

### Synthetic tandem domains of A3C have increased antiviral activity and are better packaged into virions

Of the four antiviral A3s, three of these, A3D, A3F, and A3G, all have two cytidine deaminase domains, whereas A3C has one cytidine deaminase domain and only weakly inhibits HIV-1 (17, 24, 25). These observations led us to hypothesize that synthetic double domain variants of the weakly active A3C would be more active than the single domain counterparts. To determine the antiviral effects of single versus double domain A3s, we created synthetic tandem domain A3Cs, called A3C-A3C here, using both A3C_S188,_ the variant that is most common in the human population, and A3C_I188_, the variant that has increased antiviral activity and is present at a low frequency in humans (17).

We modeled the synthetic tandem deaminase protein after the most closely sequence-related double deaminase *A3* genes, *A3D* and *A3F*. A3C*-*A3C constructs were created by fusing two A3C sequences connected with a linker sequence (Arg-Asn-Pro) that is naturally found between the N- and C-terminal domains of A3D and A3F, as well as the natural deletion in the N-terminus of the second domain of A3C such that the C-terminal domain of A3C*-*A3C begins at the fifth amino acid of A3C, a methionine (see Supplemental Fig. 1).

**Figure 1.**
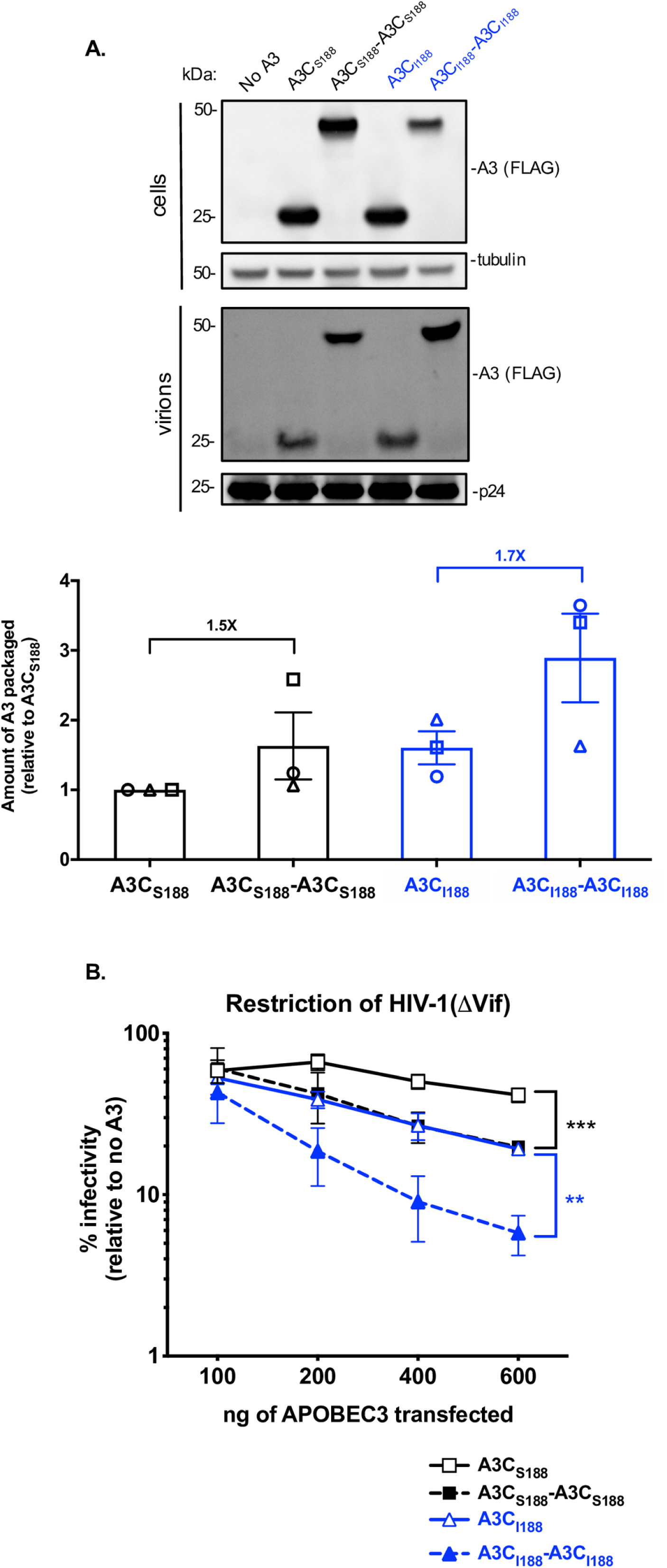
A3C tandem domain proteins have increased antiviral activity and are packaged more than their single domain counterparts. (A) Intracellular expression and virion packaging of A3C variants. HIV-1ΔEnvΔVif provirus was co-transfected into 293T cells with each A3C variant. Each variant is epitope-tagged with a 3X-FLAG tag. Top: western blot of cellular lysates probed with anti-FLAG antibody showing intracellular expression levels for A3s and tubulin as a loading control. Middle: western blot of proteins in the pelleted virions and probed with anti-FLAG antibody for A3 levels and anti-p24^gag^ for normalization. An empty vector condition was used as a negative control and labeled “No A3.” Western blot shown is representative of three biological replicates. Bottom: Bar graph showing quantification of A3 protein packaged into virions from three biological replicates (each represented by a different shape (circle, triangle, and square). A3 packaged was calculated by the abundance of A3 in the virions normalized to p24^gag^, divided by the A3 expression in the cell normalized to tubulin. The amount of A3 packaged reported is relative to A3C_S188_, which is set to 1. The fold increase in packaging of the double domain A3C relative to the respective A3C single domain is noted above the bar graph. Error bars represents standard error of the mean of the three biological replicates (+/- SEM). (B) The % infectivity of HIV-1ΔEnvΔVif pseudotyped with VSV-g and the indicated amounts of transfected plasmids of A3C variants is plotted relative to a no A3 control. A3C_S188_ (black open squares, solid line), A3C_S188_-A3C_S188_ (black closed squares, dotted line), A3C_I188_ (blue open triangles, solid blue line), and A3C_I188_-A3C_I188_ (blue close triangles, dashed blue line) are compared. Dotted lines denote the double domain A3s, while the solid lines denote single domain A3s. Data points are the mean of three biological replicates, with each biological replicate consisting of triplicate infections. Error bars show the SEM. Statistical differences were determined by unpaired *t* tests between A3C_S188_ and A3C_S188_-A3C_S188_ (black bracket) and A3C_I188_ and A3C_I188_-A3C_I188_ (blue bracket): ** P≤0.01, *** P≤0.001.

We first examined the ability of each epitope-tagged variant to be expressed in cells and packaged into virions in 293T cells. All A3C variants were expressed at similar levels in cells (Fig. 1A, top), albeit with somewhat lower expression for the A3C_I188_-A3C_I188_ protein. However, the tandem domain A3C proteins were better packaged into virions compared to the natural single domain A3 proteins. In virions, we find that there is about a 1.6-fold increase in packaging for A3C_S188_-A3C_S188_ compared to A3C _S188_ and about a 2.9-fold increase in packaging of A3C_I188_-A3C_I188_ relative to A3C _S188_ (quantified in Fig. 1A bottom, based on three independent experiments).

To evaluate the antiviral activity of these tandem domain proteins, we tested each variant at increasing concentrations in a single-cycle HIV-1ΔVif infectivity assay. As previously shown (17), A3C_S188_ inhibits HIV-1ΔVif only weakly, and A3C_I188_ has increased antiviral activity relative to A3C_S188_ (Fig. 1B: approximately 2-fold greater activity at the highest plasmid dosage). We found that A3C_S188_-A3C_S188_ has approximately 2-fold increased antiviral activity at every concentration relative to the A3C_S188_ single domain protein (black in Fig. 1B). These levels of enhanced antiviral activity are similar to the previously characterized A3C_I188_ (17). A3C_I188_-A3C_I188_, however, gained 3-fold greater antiviral activity when compared to its single domain counterpart A3C_I188_, restricting HIV-1ΔVif to 5.8% infectivity at its highest concentration (blue in Fig. 1B). Thus, these data show that A3C tandem domain proteins have increased antiviral activity relative to their single domain counterparts. This increased antiviral activity of the tandem domain variants relative to the parent single domains (Fig. 1B) largely correlates with the increased packaging of the tandem domain A3Cs related to the single domain A3Cs (Fig. 1A).

### Mutation of one active site in A3C_S188_-A3C_S188_ increases antiviral activity even further

A3C_S188_-A3C _S188_ contains two cytidine deaminase motifs within the A3 conserved amino acid sequence His-X-Glu-X_23-28_-Cys-Pro-X_2-4_-Cys (Supplemental Fig. 1). In A3F and A3G, the C-terminal catalytic domain exerts the key enzymatic activity, while the N-terminal catalytic domain mediates packaging into virions (7, 22, 23). Therefore, we next asked if the increase in antiviral activity of A3C_S188_-A3C_S188_ is due to two active cytidine deaminase sites and whether or not the enzymatic functions of A3C are important for restriction. We created active site point mutations by changing the essential glutamic acid to an alanine in each domain, E68A and E254A, respectively, and found that these changes did not affect protein expression levels in 293T cells (Fig. 2A). As expected, when the glutamic acid was mutated to an alanine in the single domain A3C_S188_ (called A3C_S188_ E68A), the protein completely lost its relatively weak antiviral activity (Fig. 2B). In contrast and unexpectedly, when the C-terminal active site of A3C_S188_-A3C_S188_ was mutated, A3C_S188_-A3C_S188_ E254A, the antiviral activity instead *increased* by 5.8-fold (Fig. 2B). On the other hand, when the N-terminal active site is inactivated in A3C_S188_-A3C_S188_ (called A3C_S188_-A3C_S188_ E68A), the antiviral activity did not significantly change (Fig. 2B), suggesting that this site is not necessary for the antiviral activity, but that the active site at position 254 inhibits antiviral activity.

**Figure 2.**
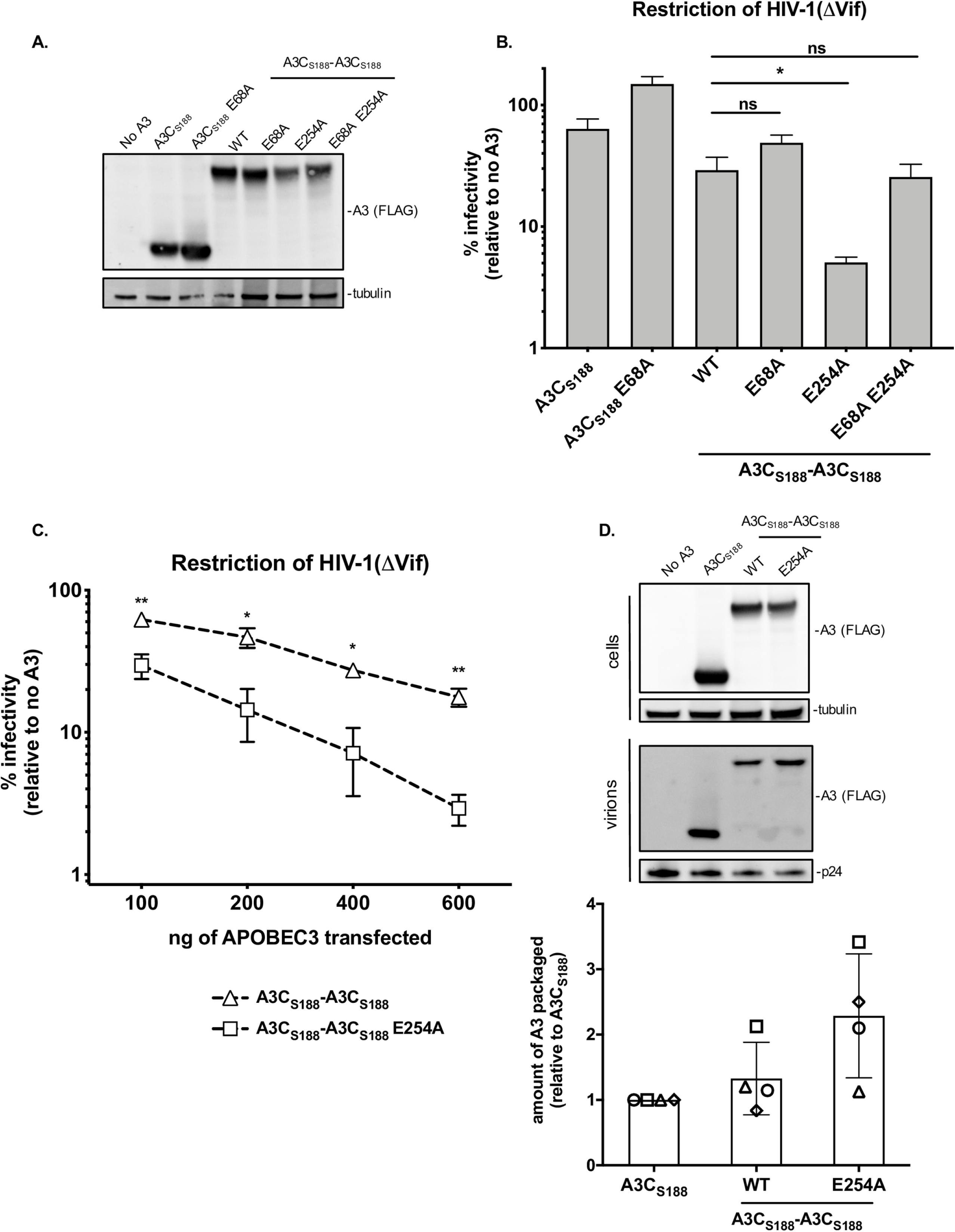
One functional deaminase domain in a tandem domain A3C protein is optimal for antiviral activity and packaging into virions. (A) Western blot analysis of intracellular expression levels of active site point mutation in A3C_S188_-A3C_S188_. WT denotes wild-type A3C_S188_-A3C_S188_; E68A refers to a mutation in the catalytic deaminase site in either the single domain A3C or N-terminus of the synthetic tandem domain A3C_S188_-A3C_S188_; E254A refers to a mutation in the C-terminus of the synthetic tandem domain A3C_S188_-A3C_S188_; E68A E254A refers to a double catalytic deaminase site mutant in A3C_S188_-A3C_S188_. Antibodies to FLAG were used to detect A3s and tubulin was used as a loading control. (B) Single-cycle infectivity assay measuring the percent infectivity of each A3C variant and active site mutant against HIV-1ΔEnvΔVif. Results from each experiment were normalized to a no A3 control. Bar graph shows the mean of three biological replicates, each with triplicate infections. Error bars represent the SEM. Statistical differences were determined by unpaired *t* tests: * P≤0.05, ns = not significant. (C) Dose response analysis showing the infectivity of HIV-1ΔEnvΔVif in the presence of increasing amounts of transfected plasmids encoding A3C_S188_-A3C_S188_ (open triangles, dotted line) compared to A3C_S188_-A3C_S188_ E254A (open square, dotted line). Infection was normalized to a no A3 control. Data points represent the mean of three biological replicates, with triplicate infections. Error bars represent SEM. Statistical differences were determined by unpaired *t* tests: * P≤0.05, ** P≤0.01. (D) Left: Representative western blot of the packaging of A3C_S188_-A3C_S188_ E254A into virions. HIV-1ΔEnvΔVif provirus was co-transfected into 293T cells with A3C variants. Cells: Intracellular expression levels western blot probed with anti-FLAG antibody for A3 levels and anti-tubulin as a loading control. Virions: Proteins in the pelleted virions shown in a western blot and probed with anti-FLAG antibody for A3 levels and anti-p24^gag^ for normalization. No A3 was used a transfection control. Right: quantification of A3 protein packaged into virions from four biological replicates. A3 packaged was calculated by the abundance of A3 in the virions normalized to p24^gag^, divided by the A3 expression in the cell normalized to tubulin. The amount of A3 packaged reported is relative to A3C_S188_, which is set to 1. Each plotted shape (circle, square, triangle, diamond) represents the A3 quantification from one biological replicate and error bars represent SEM.

We further tested the abilities of A3C_S188_-A3C_S188_ E254A and A3C_S188_-A3C_S188_ to inhibit HIV-1ΔVif in a dose-response experiment by increasing the amount of *A3* plasmid. At the highest concentrations of plasmid transfected, A3C_S188_-A3C_S188_ E254A was able to reduce the infectivity to 3%, while the wild-type A3C_S188_-A3C_S188_ only reduced infectivity to 18% (Fig. 2C). This potent antiviral activity of the A3C_S188_-A3C_S188_ E254A mutant suggests that having only one functional active site increases its antiviral effect.

In order to test if the increase in antiviral activity of the A3C_S188_-A3C_S188_ E254A mutant is also linked to increased packaging into virions, we examined the amount of packaged A3 via western blot. When compared to the intracellular expression, A3C_S188_-A3C_S188_ and A3C_S188_-A3C_S188_ E254A are expressed at relatively equal amounts (Fig. 2D). However, when we evaluate the A3 packaged into virions, we see a 2.3-fold increase in A3C_S188_-A3C_S188_ E254A packaging compared to A3C_S188_ (Fig. 2D). This finding suggests that having only one, rather than two, functional catalytic sites increases the packaging and antiviral activity of A3C_S188_-A3C_S188_.

Both cytidine deaminase-dependent and cytidine deaminase-independent mechanisms of A3 inhibition of HIV-1 have been described (1, 7–11). The antiviral activity of A3C_S188_ requires an intact deaminase motif, as demonstrated by the total loss of restriction in A3C_S188_ E68A (Fig. 2B). However, when both active sites are mutated to make a catalytically inactive A3C_S188_-A3C_S188_, we still see 3.8-fold restriction of HIV-1ΔVif – a level of activity indistinguishable from the wild-type A3C_S188_-A3C_S188_ (Fig. 2B). This suggests that the synthetic tandem domain A3C_S188_-A3C_S188_ functions as an antiviral protein in a cytidine deaminase-independent manner.

We also asked if A3C_I188_-A3C_I188_ increased antiviral activity with only one active deaminase domain. Therefore, we mutated each of the essential glutamic acids to an alanine in A3C_I188_-A3C_I188_ and tested for their ability to restrict HIV-1ΔVif. All of these A3C_I188_-A3C_I188_ active site mutants are expressed to similar levels in 293T cells (Supplemental Fig. 2A). When the C-terminal active site was mutated, A3C_I188_-A3C_I188_ E254A lost antiviral activity (2.2-fold restriction) compared to the wild-type A3C_I188_-A3C_I188_. Similar to A3C_S188_-A3C_S188_, the N-terminal active site mutant had antiviral activity that was not statistically significant from the wild-type A3C_I188_-A3C_I188_ (Supplemental Fig. 2B). However, in contrast to A3C_S188_-A3C_S188_, the C-terminally inactive mutant A3C_I188_-A3C_I188_ E254A and the double inactive mutant A3C_I188_-A3C_I188_ E68A E254A were indistinguishable in their antiviral activity, suggesting that A3C_I188_-A3C_I188_ only uses one catalytic site, like A3F and A3G. Thus, we speculate that A3C_I188_-A3C_I188_ is already optimized for its most potent restriction.

### The A3C synthetic tandem domain antiviral activity does not correlate with hypermutation activity

In order to directly test the ability of the A3C-A3C variants to hypermutate HIV-1 ssDNA, we developed an assay to assess G-to-A mutations by sequencing large numbers of unintegrated viral DNA products 12 hours post infection. This assay utilizes primers with unique barcodes designed to distinguish between A3 mutations and Illumina sequencing error (26). A plasmid-only control was used to identify sequencing error rates and the frequency of mutations acquired during PCR. A “No A3” control was used to distinguish mutations made by reverse transcriptase from those introduced by the A3 variants and any residual A3 activity in 293T cells. Each sequencing read spans the same length of HIV-1 *pol*. The number of unique sequencing reads for each condition ranged from 90,833 to 905,916. Only 0.2% of the reads from the plasmid-only control have any G-to-A mutations (Fig. 3A). 7% of the reads from the infection without any added A3 have a G-to-A mutation (Fig. 3B), presumably the result of mutations from reverse transcriptase or residual A3 activity from 293T cells. In contrast, A3G shows the highest rate of G-to-A hypermutation with 88% of reads that have at least one mutation and with over 30% of the reads having 10 or more mutations. A3G also induces a distribution of templates having 2 to 9 of such mutations (Fig. 3C). In contrast, A3C_S188_ causes fewer G-to-A mutations than A3G with 27% of reads containing at least one mutation and 9% having two or more mutations, as shown with the shift towards the left in the bar graphs relative to A3G (comparing Fig. 3C to Fig. 3D). As expected from the single-cycle infectivity assay (Fig. 2B), the mutational burden induced by the A3C_S188_ E68A active site mutant is not above background levels – only 5% of reads have any G-to-A mutations and 2% of reads have two or more of such mutations (Fig. 3E).

**Figure 3.**
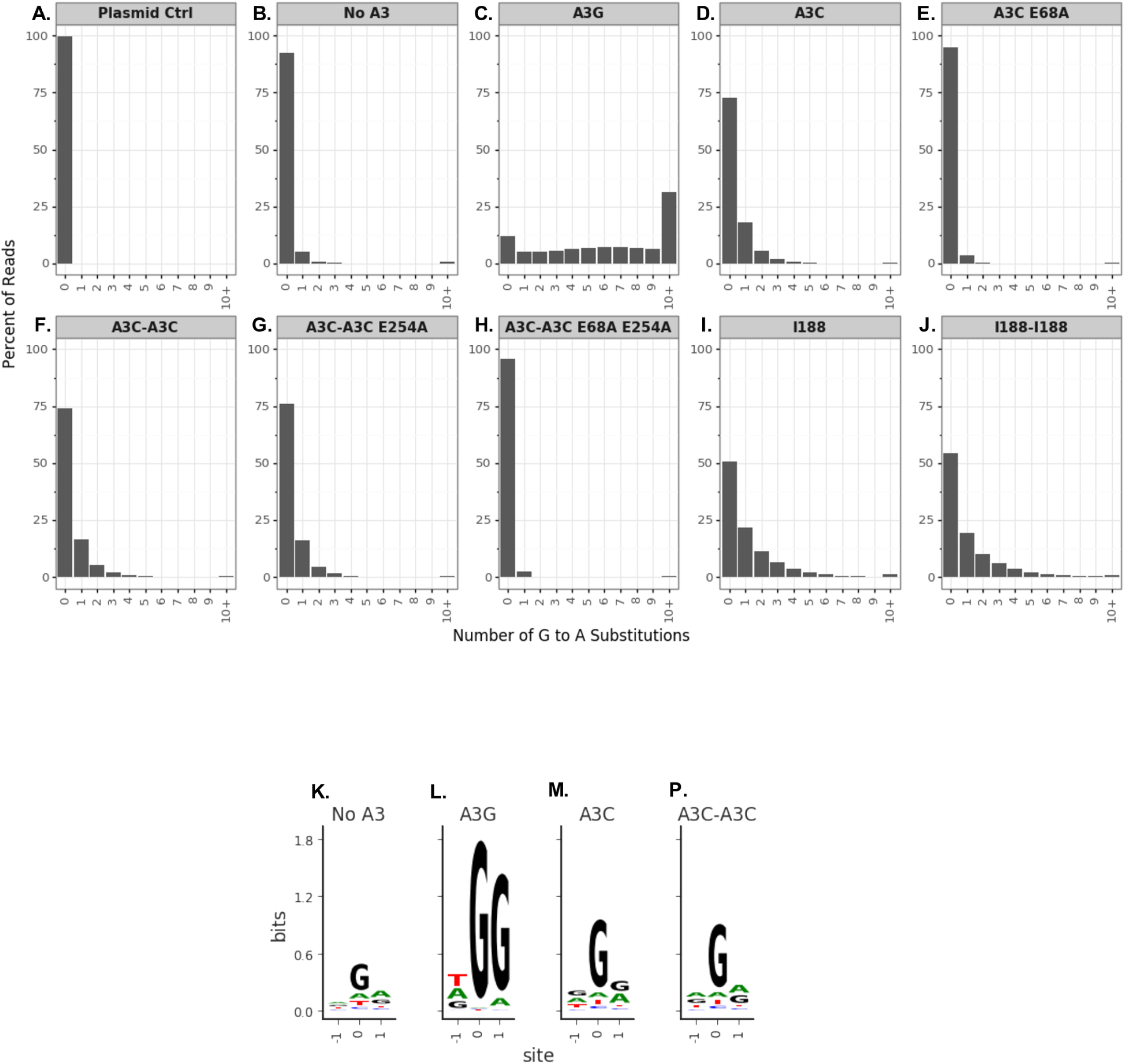
Percent of G-to-A mutations does not increase in A3C tandem domain proteins. (A-J) SUPT1 cells are infected with HIV-1 virus containing A3 and harvested for viral cDNA 12 hours post-infection. Paired-end sequencing reads were analyzed for G-to-A mutations in a region of *pol*. A plasmid control in the absence of an infection (A) is used as a sequencing control and a no A3 (B) sample was used to distinguish background mutations, including reverse transcriptase mutations. Data is shown as bar graphs of the percent of reads by the number of G-to-A substitutions in each read for A3 tested: A3G (C), A3C_S188_ (D), A3C_S188_ E68A (E), A3C_S188_-A3C_S188_ (F), A3C_S188_-A3C_S188_ E254A (G), A3C_S188_-A3C_S188_ E68A E254A (H), A3C_I188_ (I), A3C_I188_-A3C_I188_ (J). G-to-A mutations are calculated as the average frequency of G-to-A mutations between the technical replicates and read counts are shown as the sum of the reads for both replicates. (K-N) Extracted sequence information for all mutations (and neighboring bases) in the No A3 (K), A3G (L), A3C_S188_ (M), or A3C_S188_-A3C_S188_ (N) samples on viral cDNA. The constructed three-nucleotide logo plots are centered on the site of substitution, such that site 0 is fixed as the site of substitution and the -1 and +1 sites indicate the nucleotides immediately 5’ and 3’, respectively, of any substitution in these samples. The letter height, measured in bits representing information content, shows the enrichment of that nucleotide in the sample compared to the plasmid control.

Consistent with the result that much of the antiviral activity of A3C_S188_-A3C_S188_ does not depend on an active cytidine deaminase (Fig. 2B), we find that A3C_S188_-A3C_S188_ does not increase G-to-A mutations when compared to A3C_S188_ (compare Fig. 3F and 3D). For the A3C_S188_-A3C_S188_ and A3C_S188_ samples, 26% and 27% of reads have any G-to-A mutations, respectively, and the distribution of the G-to-A mutation plots are nearly identical. Interestingly, 24% of reads from the A3C_S188_-A3C_S188_ E254A sample have a G-to-A mutation, resulting in a similar to distribution as the bar graph for A3C_S188_-A3C_S188_ (Fig. 3G and 3F, respectively). This finding complements the increase in packaging results seen in Fig. 2D. As an additional control, A3C_S188_-A3C_S188_ E68A E254A has a G-to-A substitution rate below background levels (Fig. 3H).

Previous studies have shown that the increase in antiviral activity of A3C_I188_ is due to an increase in enzymatic activity (17). This finding is replicated here with a shift towards the right in the A3C_I188_ bar graph when compared to A3C_S188_ (comparing Fig. 3I and 3D) such that 49% of reads have at least one G-to-A mutation. A3C_I188_-A3C_I188_, despite having more antiviral activity than A3C_I188_ (Fig. 1A), does not have a higher percentage of mutations—49% of reads have at least one mutation in A3C_I188_ and 45% in A3C_I188_-A3C_I188_ (Fig. 3I and 3J). In sum, the hypermutation activities for the double domain A3Cs appear to have nearly identical distributions as the corresponding single domain A3Cs. Taken together, these deep sequencing data of A3 hypermutation support our earlier conclusion from the double active site mutations (Fig. 2B) that the increase in antiviral activity is not due to an increase in enzymatic activity.

In addition, we determined if the nucleotide preferences for G-to-A mutations were changed by adding a second cytidine deaminase domain to A3C. As expected (27), A3G has a strong preference of 5’-G<u>G</u>-3’ on the positive sense strand with the -1 site having equal preference among the nucleotides present (Fig. 3K). However, both A3C_S188_ and A3C_S188_-A3C_S188_ show a similar lack of preference for the -1 and +1 position (Fig. 3M and 3N), further supporting that an increase in mutation frequency does not explain the different antiviral activities of A3C_S188_ and A3C_S188_-A3C_S188_.

### A3C_S188_-A3C_S188_ tandem domain variants reduce the accumulation of reverse transcription products

Since we did not see an increase in G-to-A mutations in the A3C-A3C variants relative to A3C alone, we suspected that, similar to studies with an A3G active site mutant (9–11), the A3C-A3C tandem domain proteins might decrease reverse transcriptase products independent of hypermutation. To test this hypothesis, we used qPCR to quantify late reverse transcription (RT) products from unintegrated viral DNA. Virus produced in the presence of A3G showed a significant decrease in relative late RT products compared to the No A3 control, while A3C_S188_ had equivalent levels of late RT products as the No A3 control (Fig. 4). However, the amount of RT products produced from virus made in the presence of A3C_S188_-A3C_S188_ was significantly reduced (2.5-fold less) relative to that made in the presence of A3C_S188_ (Fig. 4). A3C_S188_-A3C_S188_ E254A had equivalent late RT products as A3C_S188_-A3C_S188_, further confirming that the increase in antiviral activity comes from the increase in packaging into virions (Fig. 4, 2C, and 2D). Lastly, there was no significant difference in the late RT products of A3C_S188_-A3C_S188_ and A3C_S188_-A3C_S188_ E68A E254A, suggesting that inhibition of RT is likely the mechanism by which these variants act (Fig. 4 and 2B). These findings further support the hypothesis that A3C super restriction factors function through a distinct mechanism to restrict HIV-1 compared to A3C single domain proteins.

**Figure 4.**
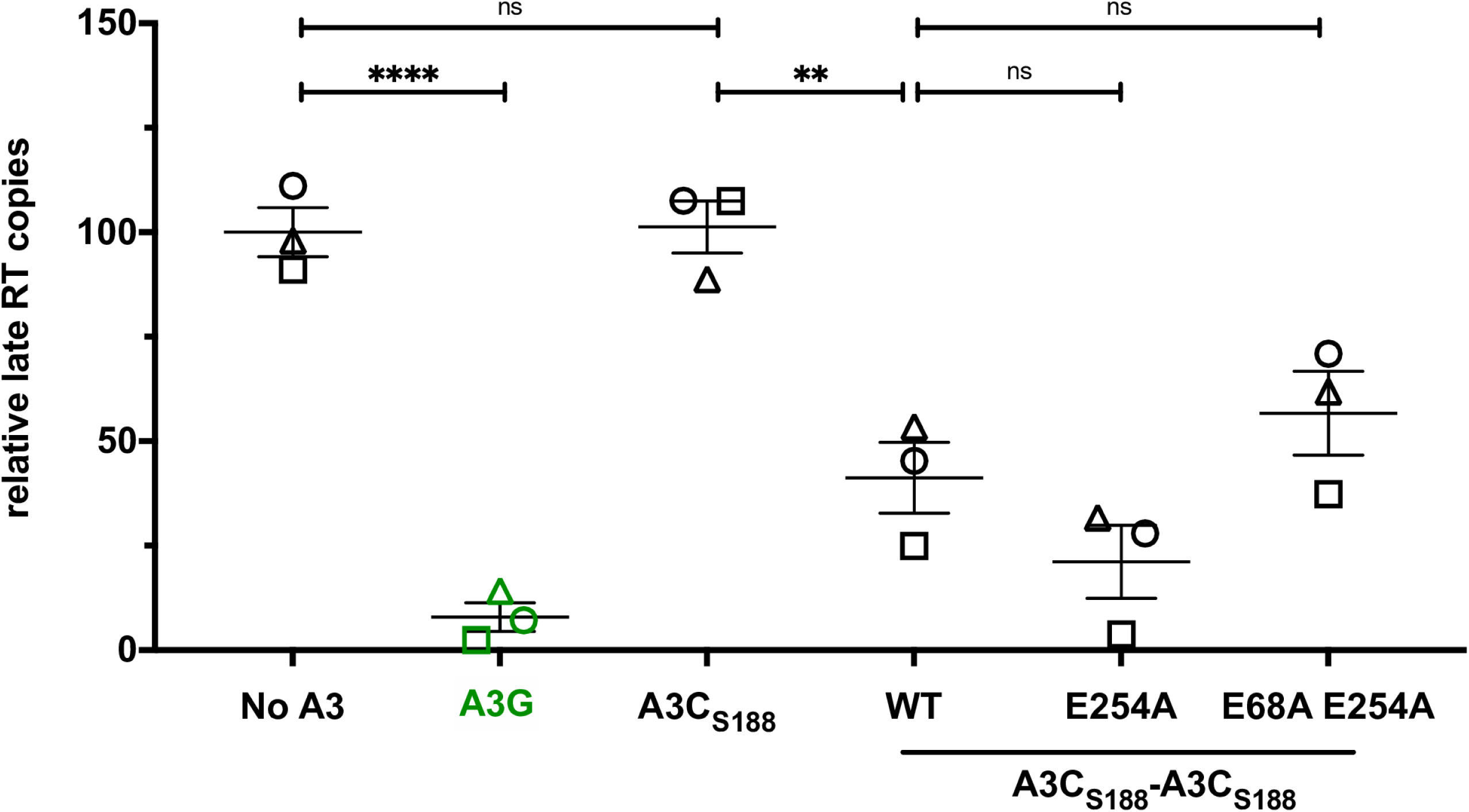
A3C tandem domain proteins operate in a deaminase-independent mechanism to inhibit reverse transcription products. (A) Copies of late reverse transcription products after infection relative to an infection of virus produced with no A3 (set to 100). SUPT1 cells were infected with HIV-1ΔEnvΔVif and either no A3 or A3 to test for inhibition of HIV-1 reverse transcription. 18 hours later, viral cDNA was harvested and the levels of HIV-1 late reverse transcription products were assayed by qPCR. The experiment was done with three separate biological replicates with each shape (circle, triangle, square), representing a normalized mean of qPCR technical duplicates for the respective biological replicate, with qPCR technical duplicates. Each sample has been adjusted for equal viral infection and a nevirapine control to subtract out DNA carry-over. Error bars indicate the SEM from three biological replicates. Statistical differences were determined by unpaired *t* tests: ** P≤0.01, **** P≤0.0001, ns= not significant.

### Tandem domain variants of A3C form larger higher-order complexes in cells relative to their native single domains

The ability of A3G to oligomerize in cells has been correlated with its antiviral activity because this oligomerization leads to increased packaging into virions (28, 29). The A3G oligomerization state also affects its binding to ssDNA and its catalytic activity (30). More specifically, A3G in lower molecular mass complexes has the ability to deaminate ssDNA (30, 31), while A3G residing in high molecular mass complexes is inactive, hinders rapid deamination of ssDNA, and is hypothesized to instead form a roadblock to inhibit reverse transcription (30, 31). Because we observed that the A3C-A3C variants inhibited late RT products (Fig. 4) rather than inducing hypermutation (Fig. 3), we examined the ability of each of the A3C variants to form high molecular weight complexes in a velocity sedimentation. As expected from previous reports (28, 32), A3G forms both lower and higher order complexes as demonstrated by its presence in the top and middle fractions of the a sucrose gradient (Fig. 5A). In contrast, both A3C_S188_ and A3C_I188_ were found in the top fractions of the gradient, overlapping with the GAPDH soluble control (Fig. 5B and Fig. 5D, respectively). This suggests the A3C single domain protein does not form higher-order complexes, unlike A3G. In contrast, we found that both A3C_S188_-A3C_S188_ and A3C_I188_-A3C_I188_ formed complexes that migrated farther down the sucrose gradient, similar to A3G (Fig. 5C and Fig. 5E, respectively). However, A3C_S188_-A3C_S188_ migrated even farther down the sucrose gradient than the pattern defined for A3G. Furthermore, both A3C_S188_-A3C_S188_ and A3C_I188_-A3C_I188_ have less protein in the top fractions than both A3G and their single domain counterparts (Fig. 5F). Together, these findings suggest that the A3C tandem domain variants reside primarily in larger higher-order complexes.

**Figure 5.**
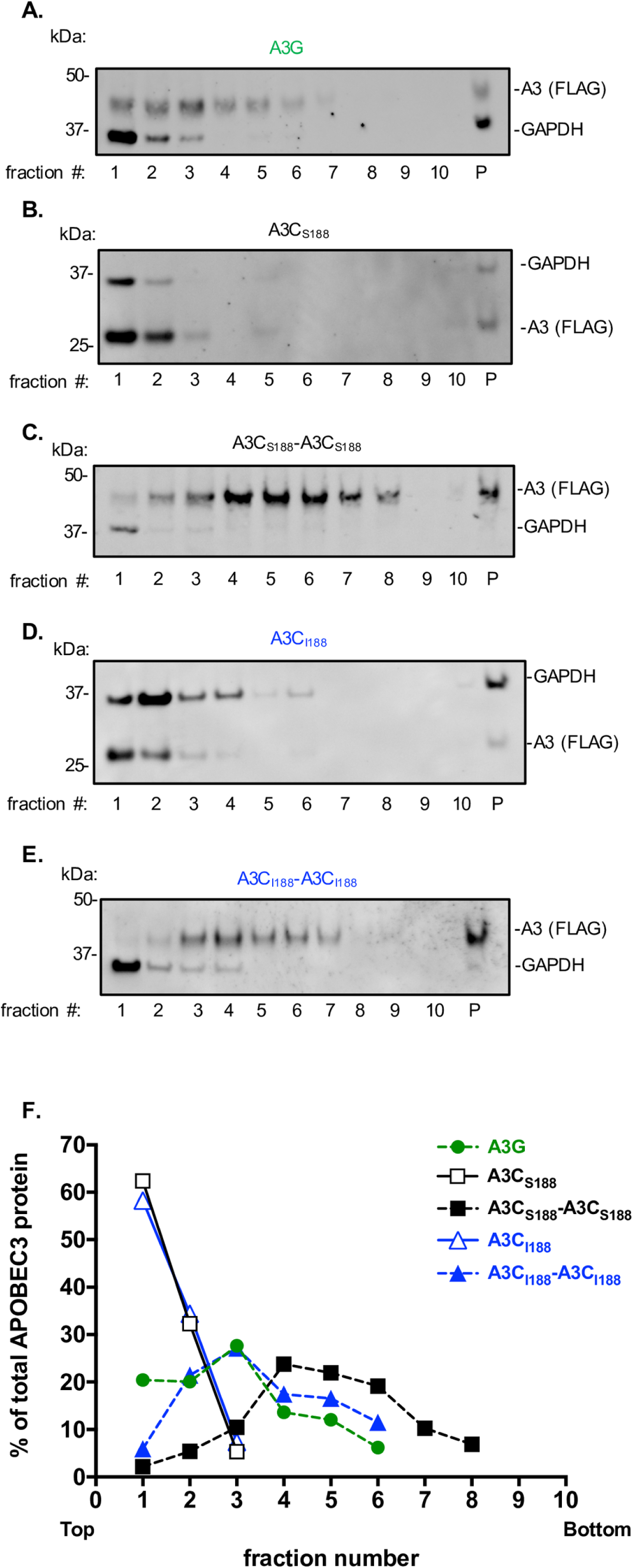
A3C double deaminase domain proteins form larger complexes than the single domains. (A-E) Velocity sedimentation and sucrose fractionation of A3C variants. Cell lysates from 293T cells transfected with A3 were subjected to velocity sedimentation fractionation (lanes 1-10, p = pellet) and 2.5% of each fraction was analyzed on a western blot: (A) A3G (B) A3C_S188_ (C) A3C_S188-S188_ (D) A3C_I188_ (E) A3C_I188-_A3C_I188_. Each western blot is probed with anti-FLAG to analyze A3s levels and anti-GAPDH to mark the soluble fraction. Higher fraction numbers are lower down the gradient and represent increasingly larger complexes. (F) Quantification of percentage of the total A3 protein in each fraction. Relative abundance of A3 in each fraction from the sucrose gradient western blots was calculated as a percentage of total APOBEC3 protein found in all the fractions combined. Dotted lines denote the double domain A3s, while the solid lines denote single domain A3s. A3G is shown in green circles, A3C_S188_ in open black squares, A3C_S188_-A3C_S188_ in closed back squares, A3C_I188_ in open blue triangles, A3C_I188_-A3C_I188_ in closed blue triangles.

### A3C tandem domain proteins are largely resistant to viral antagonism by Vif

The most effective super restriction factors would need to both increase in antiviral activity and overcome viral antagonism. Therefore, we next examined the ability of the A3C-A3C super restriction factors to escape viral antagonism. To test if HIV-1 Vif from two different strains could degrade novel A3C tandem domain protein targets, we used a single-cycle infectivity assay using a HIV provirus containing a *Vif* gene from either a lab-adapted strain (LAI) or a primary-isolate *Vif* (patient ID 1203) that has been previously shown to degrade A3H and A3C variants (33). In the absence of the viral antagonist Vif, A3G potently inhibits HIV-1ΔVif, (Fig. 6A), and, as expected, the presence of either HIV-1 LAI Vif (light purple bars) or an HIV-1 primary-isolate of Vif (dark purple bars) leads to full antagonism of A3G (Fig. 6A). While, A3C_S188_ and A3C_I188_ inhibit HIV-1ΔVif, the presence of Vif fully antagonizes this antiviral activity (Fig. 6A). In contrast, each of the A3C tandem variants is partially resistant to Vif degradation as infectivity is not completely restored (Fig. 6A). Even in the presence of Vif, A3C_S188_-A3C_S188_ still restricted HIV-1 from LAI Vif to 20% infectivity and a primary-isolate Vif to 24%. Furthermore, A3C_S188_-A3C_S188_ E254A restricted HIV-1 to 7% and 12% infectivity (LAI Vif and primary-isolate Vif, respectively), and A3C_I188_-A3C_I188_ inhibits infection to 10% with both HIV-1 Vifs (Fig. 6A).

**Figure 6.**
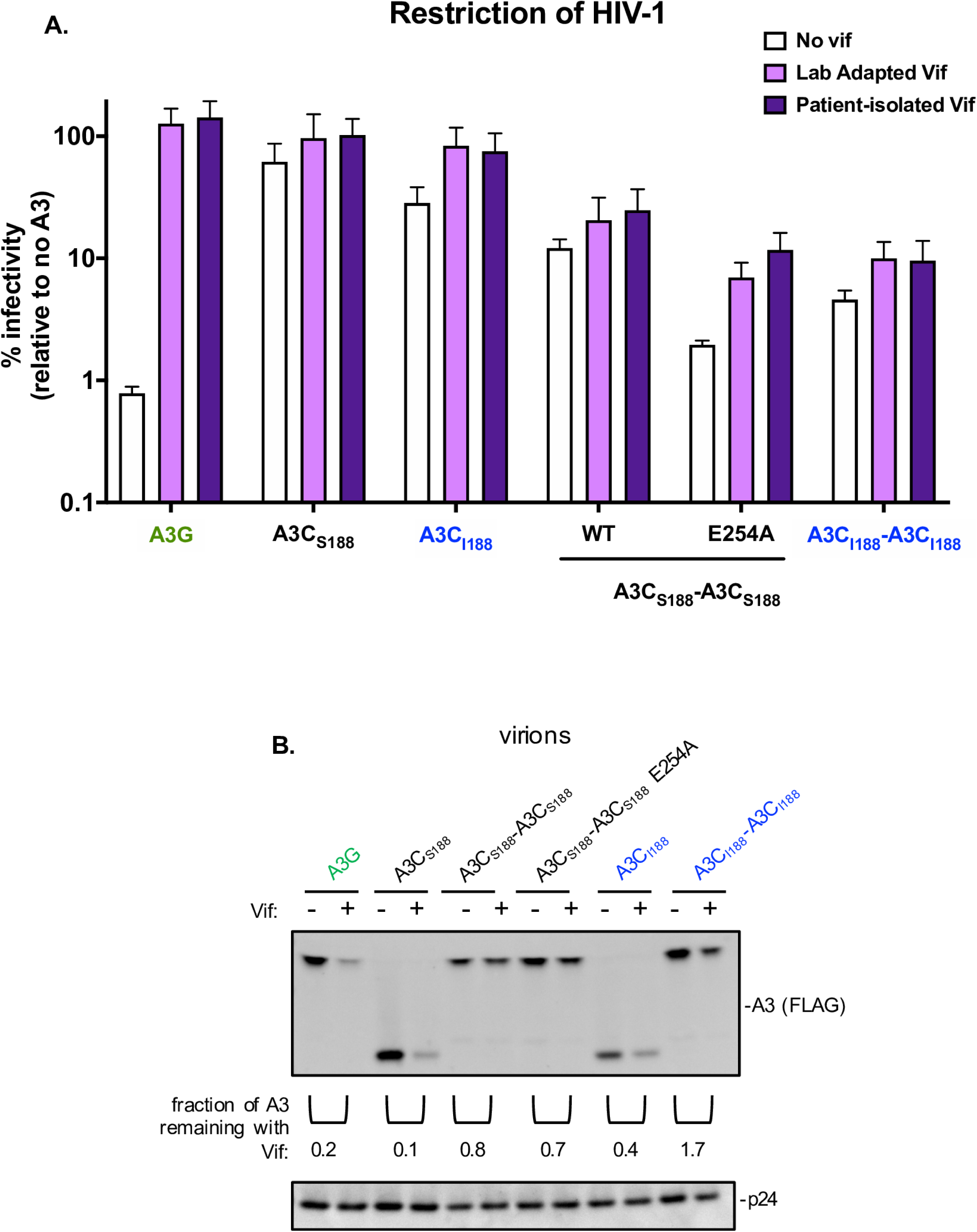
A3C tandem domain variants are resistant to viral antagonism. (A) Single-cycle infectivity assay performed in the presence of HIV-1ΔEnvΔVif (white), HIV-1ΔEnv +LAI Vif (Lab Adapted Vif, light purple), and HIV-1ΔEnv+1203 Vif (Patient-isolated Vif, dark purple). Results from each experiment were normalized to a no A3 control. Bar graph shows the mean of three biological replicates, each with triplicated infections. Error bars represent SEM. (B) Western blot of the packaging of A3C variants into virions. Either HIV-1ΔEnvΔVif or HIV-1ΔEnv+LAI Vif provirus was co-transfected into 293T cells with A3C variants. Proteins in the pelleted virions are shown in a western blot probed with anti-FLAG antibody for A3 levels and anti-p24^gag^ as a loading control. Densitometry calculations were performed to determine the fraction of A3 remaining in the presence of Vif shown under the anti-FLAG western with pairs of minus and plus Vif for each A3. Western blot shown is a representative of three biological replicates.

As expected from the inability of HIV-1 Vif to antagonize the A3C-A3C tandem domain proteins (Fig. 6A), we found that HIV-1 Vif did not decrease the amount of A3C-A3C packaged into virions (Fig. 6B). We used a western blot to evaluate levels of A3 proteins packaged into virions in the presence and absence of Vif (Fig. 6B). As a control, we used A3G, A3C_S188_, and A3C_I188_ as naturally found A3s that are degraded by HIV-1 Vif. Using densitometry, we quantified and compared the fraction of A3 probed in the presence of an HIV-1 provirus containing a deletion in the *Vif* gene to a LAI HIV-1 provirus retaining the *Vif* gene. As expected, A3G has 0.2 fraction of A3 remaining in the presence of Vif, A3C_S188_ has 0.1, and A3C_I188_ has 0.4, since HIV-1 Vif is able to degrade the naturally found A3s (Fig. 6B). Furthermore, these data support rescue of infection observed in the single-cycle infectivity assay (Fig. 6A). We found that A3C_S188_-A3C_S188_, A3C_S188_-A3C_S188_ E254A, and A3C_I188_-A3C_I188_ are packaged into budding virions in the presence of HIV-1 Vif with a fraction of A3 remaining in the presence of Vif to be 0.8, 0.7, and 1.7, respectively (Fig. 6B). This fraction of A3 remaining is much higher than the single domain counterparts and supports the inhibition of HIV-1 infection in the presence of Vif seen in the single-cycle infectivity assay (Fig. 6A and 6B). Since HIV-1 Vif from two divergent strains is unable to fully antagonize these A3C-A3C variants, our data show that a synthetic tandem domain version of an A3 protein can both increase its potency and allow it to resist antagonism by HIV-1 Vif.

## Discussion

Although human A3C can weakly inhibit HIV-1 replication, here we show that a super restriction factor can be created by linking two A3C sequences. These tandem domain proteins both increase anti-HIV-1 activity and yield a restriction factor that is partially resistant to antagonism by HIV-1 Vif. We have shown that this increase in antiviral activity is mostly explained by increased packaging of tandem domain A3s into budding virions. Moreover, we found that A3C_S188_-A3C_S188_ appears to use both a cytidine deaminase-dependent and independent mechanism for HIV-1 restriction. However, we could further increase the antiviral activity of A3C_S188_-A3C_S188_ by mutating the C-terminal active site (A3C_S188_-A3C_S188_ E254A). We found that A3C-A3C tandem domain proteins do not have an increase in mutation frequency when compared to their single domain counterparts, but rather the dominant mechanism of restriction is through inhibition of reverse transcription. Additionally, all the A3C tandem domain variants are resistant to HIV-1 Vif degradation. Together, these data point towards a selective advantage of double versus single domain A3 proteins against a viral target and support the idea that additional potency and escape from viral antagonism can be derived from combinations of APOBEC3 domains that have not been sampled in nature.

### Two domain A3C proteins are packaged at higher levels than single domain A3C proteins

In order for the A3 proteins to be antiviral, they must be packaged into virions to function during the reverse transcription process that occurs in the target cell. We found that increase in antiviral activity of the A3C tandem domain proteins increase relative to their single domain counterparts closely parallels both the increase of A3 packaged into budding virions (Fig. 1 and 2). However, the variability of the packaging assay does not allow us to conclude that packaging alone is responsible for the increased antiviral activity. Nonetheless, each of the A3C-A3C variants formed larger higher-order complexes when compared to their single domains (Fig. 5) and it has been shown that oligomerization is important for A3G to be packaged into virions (29). A3H is an interesting exception to the idea that double cytidine deaminase A3 proteins have better oligomerization characteristics relative to single cytidine deaminase A3 proteins because it is the most distantly related A3 with only one deaminase domain that never duplicated or recombined to form a double deaminase protein (5). However, similar to A3G and the A3C-A3C variants, A3H can form large complexes in sucrose gradients, yet these complexes are RNA dependent (34, 35). Also, A3H is polymorphic in human populations like A3C, however, A3H haplotype II confers strong antiviral activity (25). Recent structural work has shown that A3H binds to RNA to make functional dimers (35–37). This suggests that perhaps A3H evolved an independent mechanism to form higher-order structures that is dependent on RNA, but that A3C is unable to do this unless the second cytidine deaminase domain is artificially engineered. Furthermore, we can speculate that nature has selected for single and domain A3s because of the selective advantage of both monomer and dimer populations for deaminase-dependent activity as well as a larger order complex population for deaminase-independent activity.

### Deaminase-dependent and deaminase-independent mechanisms of super restriction factor antiviral activity

Naturally found A3 proteins primarily restrict HIV-1 through hypermutation of ssDNA intermediates during reverse transcription. Studies have reported that A3G and A3F can also function through a cytidine deaminase-independent mechanism to restrict HIV-1 (7–10). However, hypermutation rather than steric inhibition are the primary modes of restriction (7, 8, 11). Furthermore, A3G forms both low and high order complexes in cells, but the A3G residing in the higher-order complexes has been shown to have hindered deaminase activity (30). While A3G residing in the smaller complexes appears to be deaminating ssDNA targets, these large A3G complexes have been hypothesized to form a roadblock to inhibit reverse transcription (30). Interestingly, the A3C_S188_-A3C_S188_ double inactive site mutant (A3C_S188_-A3C_S188_ E68A E254A) still retained antiviral activity that, in fact, is indistinguishable from the wild type A3C_S188_-A3C_S188_ (Fig. 2B). The A3 mediated hypermutation assay data added to these findings by showing that the tandem domain A3 proteins do not have more enzymatic activity than their single domain counterparts (Fig. 3). Rather, both A3C_S188_-A3C_S188_ and A3C_S188_-A3C_S188_ E254A inhibited the total late RT products more than their A3C single domain counterparts (Fig. 4). Each of these results is also consistent with a lack of total A3 in the top fraction of the A3C tandem domains (Fig. 5). This finding supports the hypothesis that these large higher-order complex contribute to hindering reverse transcription in a deaminase-independent mechanism. Thus, we found that these A3C-A3C variants gained a new mechanism to restrict reverse transcription: the ability to inhibit HIV-1 reverse transcription, most likely through forming large complexes of oligomers.

### Only one cytidine deaminase domain is active in tandem domain APOBEC3 proteins

A3G and A3F have two deaminase domains, yet these naturally found A3s primarily rely on one domain, the C-terminal domain, for their catalytic activity (22, 23). Our finding that only one active domain is more antiviral than two active domains (Fig. 2) is similar to how the A3F and A3G domains have evolved. However, A3G and A3F do not have enhanced antiviral activity when active site mutations are created (7, 23). Additionally, the preferred enzymatic domain for A3C_S188_-A3C_S188_ is the N-terminal domain, unlike with A3F and A3G. Nevertheless, the C-terminal active site mutant of A3C_S188_-A3C_S188_ can parallel the evolutionary process that the naturally found A3s have undergone. Since A3C_S188_-A3C_S188_ is less potent as a restriction factor than A3C_S188_-A3C_S188_ E254A, this suggests that there is a disadvantage to having two fully active deaminase sites. We speculate that the specialization of the A3 domains could be important for optimal efficiency as an enzyme, such that one domain is primarily used for packaging into virions and oligomerization while the other is important for scanning ssDNA for its deamination activity. Since A3F and A3G have evolved to only use one domain for hypermutation, mutating one catalytic site will not increase antiviral activity. However, A3C_S188_-A3C_S188_ can be further improve upon by mutating the protein such that only one active site is used, A3C_S188_-A3C_S188_ E254A, to recapitulate natural selection seen in A3F and A3G.

### A3C-A3C super restriction factors are mostly resistant to Vif antagonism

Previous studies have shown that HIV-1 Vif binds and degrades A3C_S188_ and A3C_I188_ (17, 38). One viral protein, Vif, must counteract the antiviral activity of multiple A3s to achieve maximal infectivity for the virus (2). HIV-1 Vif has evolved three separate interfaces in order to degrade A3s: one for binding A3G, another for A3H, and a third interface that is able to interact with A3C/A3D/A3F (2). In contrast to the human A3 proteins, HIV-1 has never evolved to antagonize an A3C-A3C tandem domain protein. The synthetic nature of these tandem domain A3s may explain why HIV-1 Vif is not able to completely antagonize any A3C-A3C variants (Fig. 6). Since the two different strains of HIV-1 Vif are approximately 85% identical and cannot fully antagonize A3C-A3C tandem domain proteins, it suggests a potential mechanism that the interface Vif previously used to bind to the A3C is now occluded by these tandem domain proteins. However, another possibility is that A3C-A3C is packaged so efficiently that Vif is unable to target all the active A3 prior to packaging into virions, although only a ∼1.5-fold increase in packaging of the super restriction factors makes this less likely. In either case, the increased activity and resistance to Vif antagonism of these super restriction factors provide useful insights about the initial constraints of Vif recognition to novel A3 variants.

## Materials and Methods

### APOBEC3C Tandem Deaminase Domain Sequences

The A3C-A3C synthetic tandem domain was designed after the most closely related A3, A3F. The N-terminal subunit consists of amino acid residues 1-187 of A3C (residue numbers 188-190 were deleted) and was codon optimized to distinguish the N-terminus from the C-terminus. The codon-optimized N-terminus was linked to the single domain human A3C at the C-terminus using a short amino acid sequence that is found in A3F and A3D, Arg-Asn-Pro (RNP), and naturally found in A3C single domain. The C-terminal domain begins at amino acid 6 (15 nucleotide deletion) to include this RNP sequence and extends to include the remainder of A3C followed by a flexible linker and a triple FLAG tag. The A3C-A3C tandem domain was constructed using overlapping extension PCR, as described in (17) and cloned into a pcDNA4/TO vector (ThermoFisher, V102020) using BamHI/XbaI restriction sites. All point mutations were made using Site Directed Mutagenesis QuikChange II Kit (Agilent; 200524).

### Cell Culture and Transfections

HEK293T cells (ATCC, CRL-3216) were cultured in DMEM (Gibco, #11965092) with 10% HyClone Fetal Bovine Serum (GE Healthcare, #SH30910.03) and 1% penicillin-streptomycin (Gibco; #15140122) at 37°C in a humidified CO_2_ incubator. SUPT1 cells, acquired from ATCC (CRL-1942), were maintained similarly but in RPMI medium (Gibco, #11875093), 10% HyClone Fetal Bovine Serum (GE Healthcare, #SH30910.03), 1% penicillin-streptomycin (Gibco, #15140122), and 10mM HEPES. HEK293T cells were plated in 6-well dishes for transfections at a density of 1.5×10^5^ cells per 1mL. Transfections were performed with the TransIT-LT1 transfection reagent (Mirus, #MIR2304) at a reagent:plasmid DNA ratio of 3:1.

### Intracellular Protein Expression and Packaging Experiments

Intracellular expression of APOBEC3 proteins during virion production was determined by lysis of the virion-producing 293T cells with NP-40 buffer with protease inhibitor (200mM NaCl, 50mM Tris pH 7.4, 0.5% NP-40 Alternative, 1mM DTT, and Roche Complete Mini, EDTA-free tablets; 11836170001). Lysate samples were resolved on an 4-12% SDS-PAGE gel using MES buffer and transferred to a nitrocellulose membrane for western blot analysis. Anti-FLAG (Sigma; F3164), antitubulin (Sigma; T6199), and anti-p24gag (NIH-ARP; 3537) (39, 40) antibodies were used for western blots at a dilution of 1:5,000. StarBright Blue 700 Goat Anti-Mouse IgG (BIO-RAD, 12005866) were used to detect primary antibodies at a dilution of 1:5,000. The chemiluminescent signals from all western blots were imaged using a ChemiDocMPImaging System (Bio-Rad) and images were processed using Fiji/ImageJ software to quantify the densitometry for each antibody detected band.

The amount of A3 packaged into virions was evaluated by co-transfecting 600ng pcDNA4/TO.A3.3XFLAG and 1000ng HIV-1ΔVifΔEnvLuc2 (unless denoted) in a 6-well plate. Three days post-transfection, cell lysates were harvested as described above and 1.5mL of the supernatant was collected, filtered through a 0.2 micron filter, and spun down using a tabletop microcentrifuge for 1 hour at max speed at 4°C to pellet the virions. The supernatant was aspirated off and 25uL of NuPAGE 4X loading dye (Invitrogen, #NP0007) was added to each sample. Samples were boiled for 10min at 95°C and loaded on an SDS-PAGE gel.

### Single-cycle Infectivity Assay

Single-cycle infectivity assays using HIV-1 were previously described (33). 293T cells were seeded at a density of 1.5×10^5^ cells/mL in a 6-well plate. The following day, cells were transfected with 600ng provirus (HIV-1ΔVifΔEnvLuc2, unless otherwise noted), 100ng L-VSV-G, and 400ng pcDNA4/TO.A3.3XFLAG or pcDNA4/TOPO empty vector unless otherwise indicated. 72 hours later, virus was harvested and normalized for virion production using an RT-qPCR assay, as described (41, 42). A volume of virus equivalent to 2000mU/mL of RT were used for infection of SUPT1 cells. For infectivity assays, SUPT1 cells were seeded at 2×10^4^ cells per well in a 96-well plate in media containing 20 ug/mL DEAE-dextran. 72 hours later, infected cells were lysed in luciferase lysis reagent (Bright-Glo, Promega #E2610) and luciferase expression was measured on a luminometer (LUMISTAR Omega, BMG Labtech). Infectivity of each virus was normalized to 100% based on a No A3 control. All HIV-1 constructs are based on the LAI strain. A clade B patient derived Vif (PID:1203) that is able to antagonize A3H haplotype II was previously described (33).

### Deep Sequencing of A3 Mediated Mutations

To analyze A3 mediated mutations, 293T cells were seeded at a density of 1.5×10^5^ cells/mL in a 6-well plate. The following day, cells were transfected with 600ng provirus (HIV-1ΔVifΔEnvLuc2), 100ng L-VSV-G, and 400ng pcDNA4/TO.A3.3XFLAG or pcDNA4/TO empty vector. Three days later, virions were harvested and quantified similar to single-cycle infectivity assay mentioned above. 2×10^6^ SUPT1 cells were infected with 500uL of each Benzonase-treated virus with and without a 10mM nevirapine control and spinoculated at 1,100 x g for 30min at 30°C. Twelve hours later, unintegrated viral cDNA was isolated using Qiagen Miniprep Kit (QIAprep Spin Miniprep Kit, 27106).

To determine A3 mediated mutations, we used a barcoded Illumina deep sequencing approach previously described (43) and explained in more detail at https://jbloomlab.github.io/dms_tools2/bcsubamp.html. This approach attaches unique molecular identifiers (8*N) to each DNA molecule, which enables increased correction of PCR and sequencing errors by determining molecule-specific consensus sequences. A region of *pol* (44, 45) was amplified using KOD Hot Start Master Mix (Millipore, 71842) in a first round of PCR with forward primer: CTTTCCCTACACGACGCTCTTCCGATCTNNNNNNNNGACAAGGAACTGTATCCTTT AACTT and reverse primer: GGAGTTCAGACGTGTGCTCTTCCGATCTNNNNNNNNCTGGTACAGTTTCAATAGGA CTAAT. The first round PCR was carried out using the following parameters: 95°C for 2min and 20 cycles of 95°C for 20sec, 70°C for 1sec, 50°C for 10sec, 70°C for 2 min, followed by a 95°C for 1min and a hold at 4°C. PCR products were cleaned up using AMPure beads (Beckman Coulter, A63880) and quantified with Qubit dsDNA HS Assay Kit (ThermoFisher, Q32854). Next, round 1 PCR products were bottlenecked such that each uniquely barcoded ssDNA molecules would be read ∼2.9 times during sequencing. A second round of PCR performed with KOD Hot Start Master Mix (Millipore, 71842) supplemented with 1mM MgCl_2_ under the following parameters: 95°C for 2min and 23 cycles of 95°C for 20sec, 70°C for 1sec, 60°C for 10sec, 70°C for 10sec, followed by a hold at 4°C. Samples were again cleaned up, quantified, pooled, purified via gel electrophoresis, and sequenced on an Illumina MiSeq, using 2×250 paired-end reads. Deep sequencing was carried out in technical replicate for each experimental condition, starting with the initial PCR reactions.

We used the software package dms_tools2 to align sequencing reads and build consensus sequences for each uniquely tagged DNA molecule (46). Error-corrected reads were compared to the target sequence to determine the number, identity, and surrounding nucleotides of all substitutions in each read. Reads with high numbers of substitutions (>10% of non-G nucleotides) at the junction of the two paired- end reads were removed from the analysis as these substitutions were most often found to be alignment artifacts. Since A3s are known to cause G-to-A substitutions, we initially subsampled our data to look only at G-to-A substitutions. We plotted the frequency of reads in each sample with a given number of G-to-A mutations (0, 1, 2, etc. up to 9, and then 10+) (Fig. 4A-J). Frequencies were calculated as the average frequency of technical replicates and read counts are shown as the sum of reads for both replicates.

To investigate if substitutions occurred more frequently in certain nucleotide contexts, we computationally extracted all three-nucleotide sequences centered on a substitution from each sample. This gave us a count for each sample of how many times each nucleotide was immediately upstream (site -1) of a substitution, underwent a substitution (site 0), or immediately downstream of a substitution (site 1). We then calculated the frequency for each nucleotide at each of these three sites in each sample. Data for technical replicates were combined by averaging the frequencies of each nucleotide at each site across the two replicates for each sample. Next, we calculated background-corrected frequencies for each nucleotide at each site using the frequencies in plasmid control sample as our background. We based this frequency correction on the ‘type 2’ logos from Gorodkin, et al. (47).

If *q*_*ik*_ is the frequency of base *k* at site *i* in the sample and *pik*is that frequency in the background, then the background corrected frequency is:

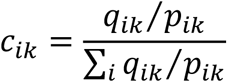

We then calculated the information content of each site as:

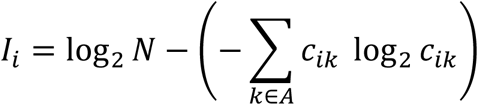

Where A is the set of nucleotides [A, C, G, T] and N is the number of elements in A (47, 48).

Finally, we calculated the letter height at each site as:

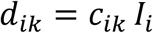

The sequencing reads were uploaded to the NCBI SRA with the BioProject accession number PRJNA605864 and run accession numbers SRR11059567 to SRR11059589. The computational pipeline to analyze the sequencing data and generate Fig. 4 is available on Github (https://github.com/jbloomlab/SuperRestrictionFactor_Hypermutation).

### qPCR Assay for HIV Late Reverse Transcription Products

For qPCR analysis of HIV late RT products, 2×10^6^ SUPT1 cells were infected with 500uL of each Benzonase-treated virus with and without a 10mM nevirapine control and spinoculated at 1,100 x g for 1hr at 30°C. 18 hours post infection, unintegrated viral cDNA was isolated using Qiagen Miniprep Kit (QIAprep Spin Miniprep Kit, 27106). HIV cDNA was amplified with TaqMan Gene Expression Master Mix (AppliedBiosystems #4369016), J1 FWD (Late RT F) – ACAAGCTAGTACCAGTTGAGCCAGATAAG, J2 REV (Late RT R) GCCGTGCGCGCTTCAGCAAGC, and LRT-P (late RT Probe) – FAM-CAGTGGCGCCCGAACAGGGA-TAMRA (49, 50). Data was acquired on an ABI QuantStudio5 Real Time (qPCR) machine and analyzed on Prism software.

### Velocity Sedimentation

For analysis of A3 migration on a sucrose gradient, 293T were harvested 72 hours post-transfection (1000ng of A3 per well in a 6-well plate) using a low salt and EDTA buffer (0.2M HEPES pH7.9, 0.1M NaCl, 0.01 MgCl_2_, 0.002 EDTA pH8.0, 3.5% triton-X 100) (51) with protease inhibitor cocktail (Roche, 11836170001). For each sample, 50uL of lysate was layered on a step gradient (10%, 15%, 40%, 50%, 60%, 70%, and 80% sucrose in lysis buffer) and subjected to velocity sedimentation in a Beckman MLS50 rotor at 45,000rpm (163,000 x g) for 37min at 4°C, as described previously (34). Gradients were fractioned (400µL) from top to bottom and analyzed via western blotting.

## Acknowledgments

We thank Nicholas Chesarino, Jaisri Lingappa, Rick McLaughlin, and Molly OhAinle for their thoughtful discussions and comments on this manuscript. The following reagent was obtained through the NIH AIDS Reagent Program, Division of AIDS, NIAID, NIH: Anti-HIV-1 p24 Monoclonal (183-H12-5C) (Cat# 3537) from Dr. Bruce Chesebro and Kathy Wehrly. Jesse Bloom is an Investigator of the Howard Hughes Medical Institute. This work was funded by UW CMB Training Grant (NIH T32GM007270) and NSF predoctoral fellowship (NSF DGE-1762114) to MM, F30 fellowship (NIAID F30AI149928) to KD, NIH/NIAID R01 AI140891 to JDB, NIH/NIAID P50AI150476 (PI: Nevan Krogan, subaward to ME), and NIH/NIAID R01 AI030927 to ME.

## Figure Legends

**Supplemental Figure 1.**

Clustal Omega amino acid alignment of A3C-A3C and the closely related A3, A3F. The “RNP” amino acid sequence (Arginine-Asparagine-Proline) that links the two deaminase domains together is highlighted in yellow. The end of the first domain and the beginning of the second domain of A3C-A3C is delineated with a red arrow. The isoleucine human polymorphism in each of the domains is shown in blue text. The conserved A3 cytidine deaminase motif, His-X-Glu-X_23-28_-Cys-Pro-X_2-4_-Cys, is highlighted in grey. The essential glutamic acid that is necessary for deaminase activity is in green text. Asterisks indicates positions which have a conserved residue between A3C-A3C and A3F; a colon denotes conservation between groups of strongly similar properties; a period indicates conservation between groups of weakly similar properties.

**Supplemental Figure 2**

Western blot analysis of intracellular expression levels of active site point mutation in A3C_I188_-A3C_I188_. WT denotes wild-type A3C_I188_-A3C_I188_; E68A refers to a mutation in the catalytic deaminase site in either the single domain A3C or N-terminus of the synthetic tandem domain A3C_I188_-A3C_I188_; E254A refers to a mutation in the C-terminus of the synthetic tandem domain A3C_I188_-A3C_I188_; E68A E254A refers to a double catalytic deaminase site mutant in A3C_I188_-A3C_I188_. Antibodies to FLAG were used to detect A3s and tubulin was used as a loading control. (B) Single-cycle infectivity assay measuring the percent infectivity of each A3C variant and active site mutant against HIV-1ΔEnvΔVif. Results from each experiment were normalized to a no A3 control. Bar graph shows the mean of three biological replicates, each with triplicate infections. Error bars represent the SEM. Statistical differences were determined by unpaired *t* tests: * P≤0.05, ** P≤0.01, ns= not significant.

